# Missing pieces in the full annual cycle of fish ecology: a systematic review of the phenology of freshwater fish research

**DOI:** 10.1101/2020.11.24.395665

**Authors:** Megan E. Brady, Andrew M. Chione, Jonathan B. Armstrong

**Affiliations:** Department of Fisheries and Wildlife, Oregon State University, Corvallis, Oregon, United States of America

**Author notes:** Corresponding author: (MEB).

## Abstract

In recent decades, fish ecologists have become increasingly aware of the need for spatially comprehensive sampling. However, a corresponding reflection on the temporal aspects of research has been lacking. We quantified the seasonal timing and extent of freshwater fish research. Since reviewing all prior work was not feasible, we considered two different subsets. First, we compiled the last 30 years of ecological research on juvenile Pacific salmon and trout (*Oncorhynchus* spp.) (n = 371 studies). In addition to the aggregate, we compared groups classified by subject matter. Next, to evaluate whether riverscape ecology has embraced space at the expense of time, we compiled research across taxa in which fish were enumerated in a spatially continuous fashion (n = 46). We found that ecological *Oncorhynchus* spp. research was biased towards summer (40% occurred during June-August) and the month of June in particular, at the expense of winter work (only 13% occurred during December-February). Riverscape studies were also biased toward summer (47% of studies) and against winter (11%). It was less common for studies to encompass multiple seasons (43% of ecological *Oncorhynchus* spp. studies and 54% of riverscape studies) and most were shorter than 4 months (73% of ecological *Oncorhynchus* spp. studies and 81% of riverscape studies). These temporal biases may cause researchers to overemphasize ecological phenomena observed during summer and limit our ability to recognize seasonal interactions such as carry-over effects or compensatory responses. Full year and winter studies likely hold valuable insights for conservation and management.

## Introduction

A key challenge in conservation is to understand how abiotic and biotic heterogeneity mediate the function of ecosystems and the survival of biota that inhabit these environments. This heterogeneity exists in both space and time, creating a shifting mosaic of physical and biological conditions that has significant ramifications for biota [1]. Phenomena ranging from ontogenetic niche shifts [2] to the stability of fisheries [3] can only be understood by jointly considering interactions between space and time. However, because resources are limited and characterizing stream heterogeneity is a non-trivial task, it is often not feasible to study multiple dimensions of variation simultaneously. Indeed, many fundamental concepts in stream ecology are either spatially or temporally focused.

For example, spatial patterns of biota are often described with minimal reference to time. This applies to early work, such as the longitudinal zonation of fishes [4], but also the River Continuum Concept [5] and the contemporary emphasis on high spatial resolution sampling found in “riverscape” ecology [6]. Although time is recognized as the “fourth dimension” of the riverscape[7] and the intersections of various temporal and spatial scales has been noted as important [6], in practice, the suffix “scape” is typically used when working at large spatial extents of data, which often compounds the challenges of incorporating time.

Similarly, time is often considered independently in studies of both habitat and fish. Stream ecologists increasingly embrace a regime approach to characterizing temporal variation in habitat conditions, originating with the Natural Flow Regime [8], which considered the statistical distribution of conditions and metrics such as event magnitude, frequency, seasonal timing, predictability, duration, and rates of change. The regime concept is now applied beyond water quantity to include aspects of water quality [9,10], as well as physical attributes such as sediment, large wood, and abundance of pools [11]. In fisheries ecology, temporal variation is probably most commonly studied in the form of population dynamics, i.e., fluctuations in abundance typically described at an annual resolution. However, many important processes that may scale up to affect population dynamics (e.g. growth) play out at intra-annual timescales and relate to seasonality.

It is often recognized that short-term datasets can be inadequate because they fail to capture historical levels of productivity (i.e. the shifting baseline) or reveal coarser scale temporal patterning such as regime shifts [12]. Likewise, for cyclically patterned temporal variation, interpretations may be misleading if they are based on a limited portion of a cycle. For example, many fish switch between habitat types throughout the diel cycle [13] so only studying animals during daytime may fail to capture important habitats. Similarly, refuge habitat identified in summer may not represent refuge habitat for other seasons and stressors [14]. Riverine systems may exhibit extreme seasonal variation, with water temperatures ranging 20°C or more and flows varying 100-fold. This strongly affects not only fish and other aquatic organisms, but also the feasibility of field sampling. While a temperature logger can effectively collect data every day of the year, the cost and logistical challenges of sampling fish vary tremendously and can strongly govern when biological data are collected. Extrapolating from data that pertain to specific points in time can lead to misleading interpretations regarding how fish behave, the production capacity for ecosystems, and what locations or habitat types are important [15,16]. This is particularly problematic in the study of mobile organisms that undergo substantial physiological and ecological changes throughout their lifetimes, such as Pacific salmonids. The objective of this paper is to characterize the temporal attributes of fish ecology research to elucidate potential data gaps and guide future research.

Recent work on birds, amphibians, reptiles, and mammals found strong seasonal biases in field research [17], but analogous work on fish has been lacking. The assertion that winter fish ecology is an important, yet understudied portion of the research portfolio is not new [18]; however, this hypothesis remains unquantified. It was not feasible to screen the research for all fish species during all life phases, so we limited our systematic review to a single taxon of fish: *Oncorhynchus* spp. We focused on juvenile Pacific salmon and trout in freshwater because they are well-studied (providing us the power to detect trends in sampling), they live in highly seasonal environments (which means an incomplete understanding of the annual cycle would be a problem and is thus important to test for), and they are distributed across multiple continents (thus representing a wide-spread species of interest). Here, we characterize the temporal aspects of freshwater fish ecological research within the taxon of Pacific salmon and trout (*Oncorhynchus* spp.) during the last 30 years. We characterized patterns in the seasonal timing and duration of ecological field studies and considered how these patterns varied across three focal topics: fish-habitat interactions, trophic ecology, and spatial distribution. This analysis of a specific taxon was then complemented with a cross-taxa analysis. We assessed whether spatially extensive sampling has come at the expense of time by reviewing the timing of “riverscape” studies across all fish taxa.

## Materials and methods

### Data screening

To determine whether and to what extent temporal biases are present in fish field research, we conducted a review of two case studies: 1) research within the *Oncorhynchus* species and 2) research across fish species within “riverscape” studies. We defined “riverscape” studies as those studies that employed the use of spatially continuous data (or nearly so) that covered a high spatial extent so that multi-scale patterns could be revealed [6] as opposed to the more typical method of using of a handful of points that are extrapolated to represent large extents. We focused on three temporal aspects of research: 1) what months and seasons juvenile salmonid ecology research occurs, 2) the duration of studies, and 3) whether seasons were studied individually or if seasonal interactions were examined.

To examine our first case study of *Oncorhynchus* research, we reviewed 13 journals that commonly publish fisheries ecology research as opposed to human consumption of fish research. Using the Web of Science database (last searched 21 August 2020), we performed the following search: TS=(salmon OR salmonids OR Oncorhynchus) AND SO=(CANADIAN JOURNAL OF FISHERIES “AND” AQUATIC SCIENCES OR Ecology OR Ecology of Freshwater Fish OR Ecosphere OR Ecosystems OR Environmental Biology of Fishes OR Freshwater Biology OR Hydrobiologia OR North American Journal of Fisheries Management OR Oecologia OR PLoS ONE OR Science OR Transactions of the American Fisheries Society) Indexes=SCI-EXPANDED, SSCI, A&HCI, ESCI Timespan=1988-2017. The past 30 years was chosen to characterize the current patterns of research. We screened the articles and selected those that dealt with the ecology of juvenile *Oncorhynchus* species during freshwater residence. The juvenile life stages of fry, parr, and smolt were all included. Only papers that presented original, ecologically focused data were included, whether they were observational studies or experimental studies conducted in a natural environment. Many studies were excluded because they were not ecological field studies. We did not include studies on fish species other than *Oncorhynchus*, laboratory studies, physiological response or manipulation research, smolt-to-adult survival studies, research dealing with the adult life stage of *Oncorhynchus* spp., studies occurring in estuarine or marine environments, studies that collected physical or biological habitat data but did not actually sample fish, reviews, or models not validated with field data. This resulted in 371 articles examined in this study (S1 Fig).

To identify temporal patterns across fish species, we identified “riverscape” studies that utilized spatially continuous sampling [6]. The term “riverscape” has been applied inconsistently but is often used to refer to sampling employing large spatial extent. We use the term to include large spatial extent sampling as well as sampling in line with the argument by Fausch et al. [6] for sampling across multiple spatial scales to observe patterns and processes playing out throughout entire river systems in order to sample rivers at the same scale that we manage them at. Using the Web of Science database (last searched 23 October 2020), we performed the following search: TS=(riverscape OR spatially continuous OR longitudinal distribution OR Fausch et al. 2002) AND TS=(fish OR fishes OR salmon) AND TS=(stream OR river OR freshwater OR lake) Indexes=SCI-EXPANDED, SSCI, A&HCI, ESCI Timespan=1988-2017. We then examined every article and selected those that dealt with spatially continuous, high spatial extent, or “riverscape”-scale sampling that included fish data collection. This resulted in 46 articles examined in this study (S2 Fig).

### Data analysis

We classified each publication for both the ecological dataset and the riverscape dataset by the temporal aspects of data collection. First, we read the Methods section of each article screened and recorded the presence/absence of data collection in each month and season. We defined seasons meteorologically as aligned with the calendar months of June 1-August 31 for summer, September 1-November 30 for autumn, December 1-February 28 for winter, and March 1-May 30 for spring. Seasons were not defined by solstice or equinox to stay consistent with presence/absence within a single month. Studies may encompass more than one month, therefore the number of data points for these analyses are greater than the number of studies included in the review. Second, we quantified the frequency of the number of meteorological seasons (1-4) that were included in these studies to analyze temporal extent and consideration of inter-seasonal interactions (i.e., carry-over effects).

To explore whether temporal aspects of sampling differed among research areas, we classified each study into three focal areas: 1) fish-habitat interactions and the impact of habitat units and types on juvenile salmonid biology or behavior, 2) trophic ecology including fish diet, foraging, and food web structure, and 3) spatial distribution including movement and landscape-scale distribution. Studies examining fish growth and survival were often presented by researchers as a function of some aspect of one of the three focal areas identified and were classified accordingly. The temporal distribution and extent of sampling effort was then quantified both collectively and by research category. Each study was only classified into one of the three focal areas based on the main objective of the study. Studies that did not fall into one of these four main categories were classified as “Other” and included in overall analysis but not the subset analyses.

### Statistical methods

We tested for temporal biases in temporal distribution and extent using Pearson _*X*_^2^-tests in R 4.0.2. Equal values would indicate that no bias exists, supporting the null hypothesis. While the test is objective, we acknowledge that the interpretation is subjective due to the assumptions that all months and seasons are equally important and present equal stresses, limitations, or opportunities for growth, fitness, and survival for juvenile salmonids.

We also acknowledge that seasonality varies with latitude, elevation, and position in watershed, so the ecological conditions associated with a particular month or season may vary among locations (and thus among the studies in our paper). Thus, the implications of the temporal biases we observed may be somewhat context dependent.

## Results

### Monthly temporal distribution of studies

At a monthly resolution across all ecological topics within juvenile *Oncorhynchus* spp. studies, we found that the most frequently represented month was 3-6 times more common than the least frequently represented month (Fig 1). December was the least represented month across all topics, while the summer months of June, July, and August were most common among topics. The month of June had a significantly higher proportion of studies than the month of December at 14% and 3%, respectively.

**Fig 1.**
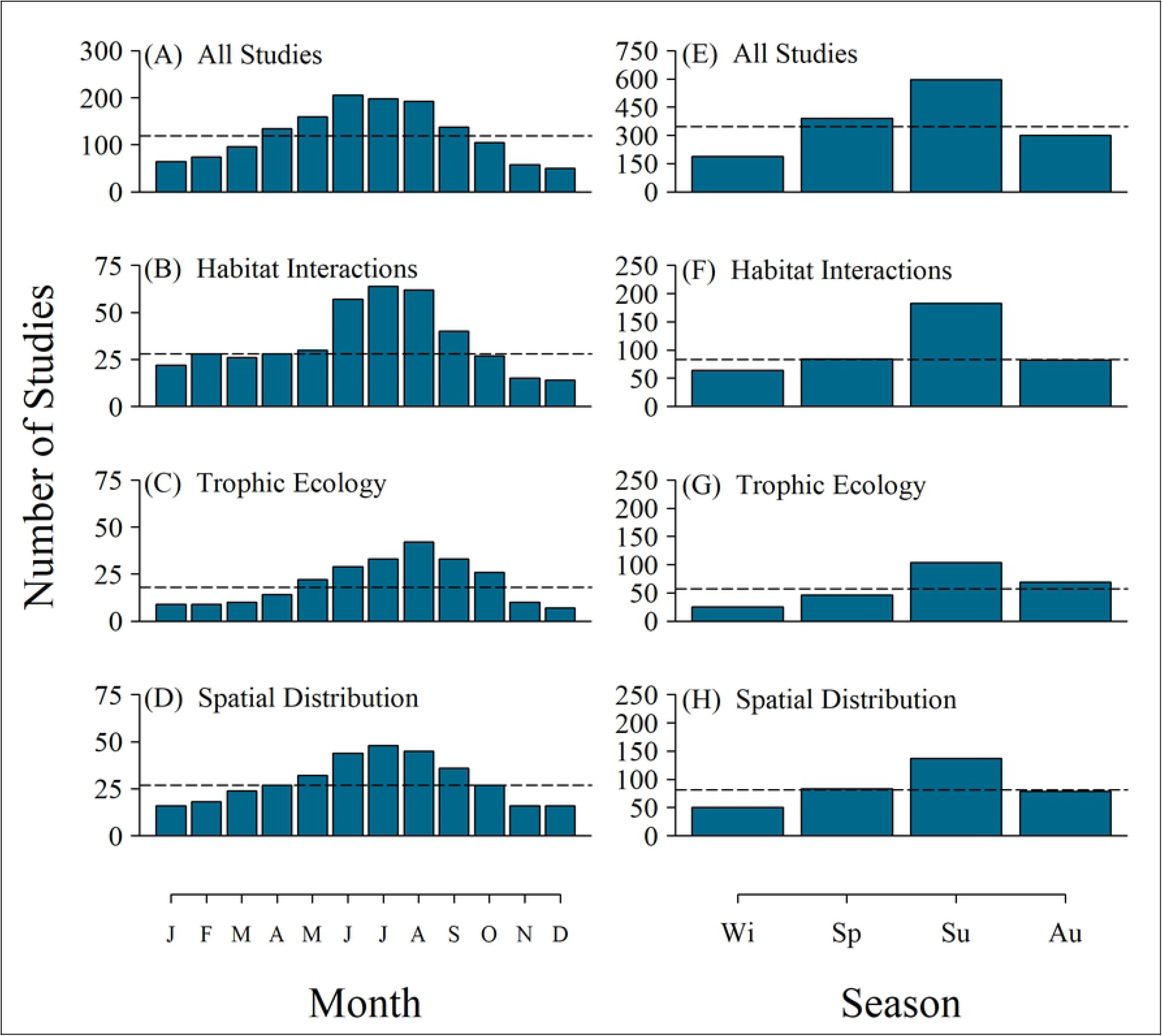
Temporal distribution of juvenile salmon ecology studies. Left column: monthly distribution (left to right: January to December) of sampling effort for juvenile Pacific salmon and trout studies from 1988-2017 for (A) all studies (X2=289.58, p < 0.0001, n=1476, median=119.5), (B) habitat studies (X2=97.421, p < 0.0001, n=413, median=28), (C) trophic ecology studies (X2=78.131, p < 0.0001, n=244, median=18), (D) spatial distribution studies (X2=53.67, p < 0.0001, n=439, median=27). Right column: seasonal distribution of sampling effort for juvenile Pacific salmon and trout studies from 1988-2017 for (E) all studies (X2=243.39, p < 0.0001, n=1476, median=345.5), (F) habitat studies (X2=84.482, p < 0.0001, n=413, median=83), (G) trophic ecology studies (X2=56.295, p < 0.0001, n=244, median=57.5), (D) spatial distribution studies (X2=45.258, p < 0.0001, n=349, median=81). The number of studies for each month or season was calculated using presence or absence of research during that time frame. Dashed horizontal lines are data median. Studies may occupy more than one month or season. Seasons were defined meteorologically, but as whole months. Summer is defined as the months June, July, and August; Autumn is defined as the months September, October, and November; Winter is defined as the months December, January, and February; Spring is defined as the months March, April, and May.

### Seasonal temporal distribution of studies

Across all ecological topics within juvenile *Oncorhynchus* spp. studies, we found that 39-44% of studies occurred during summer while only 10-15% of studies occurred during winter (Fig 1). There has been little change in the temporal distribution of research efforts with the proportion of winter studies remaining significantly lower than summer studies (Fig 2).

**Fig 2.**
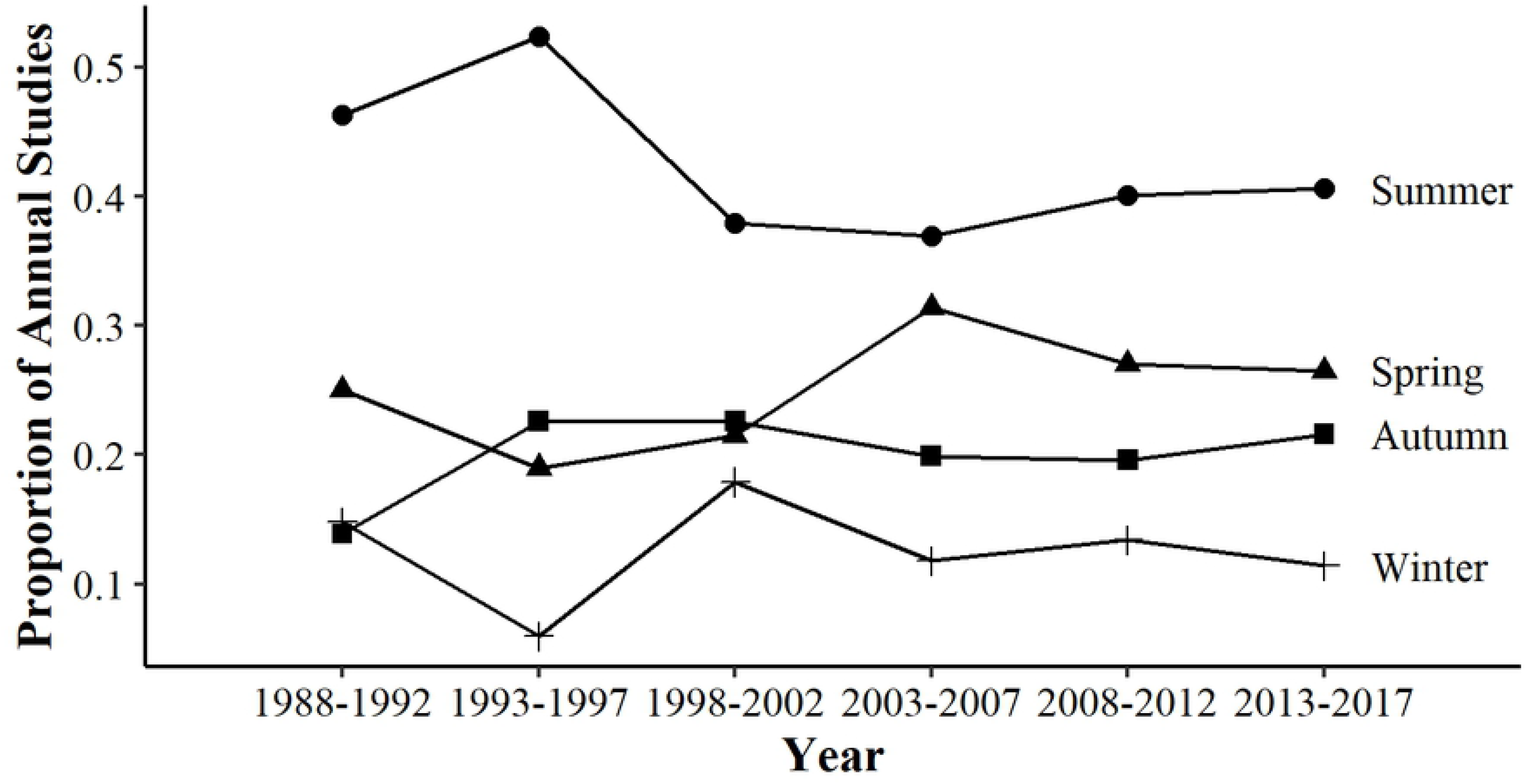
Seasonal study distribution over time. Change in the proportional temporal distribution (seasonal timing) of all studies published from 1988-2017 in 5-year increments.

### Monthly temporal extent of studies

At a monthly resolution across all ecological topics within juvenile *Oncorhynchus* spp. studies, we found that most studies had limited temporal extent across the annual cycle, with 71-75% of studies containing data from 4 months or less (Fig 3). Less than 2-8% of studies across all topics encompassed data from all 12 months of the year.

**Fig 3.**
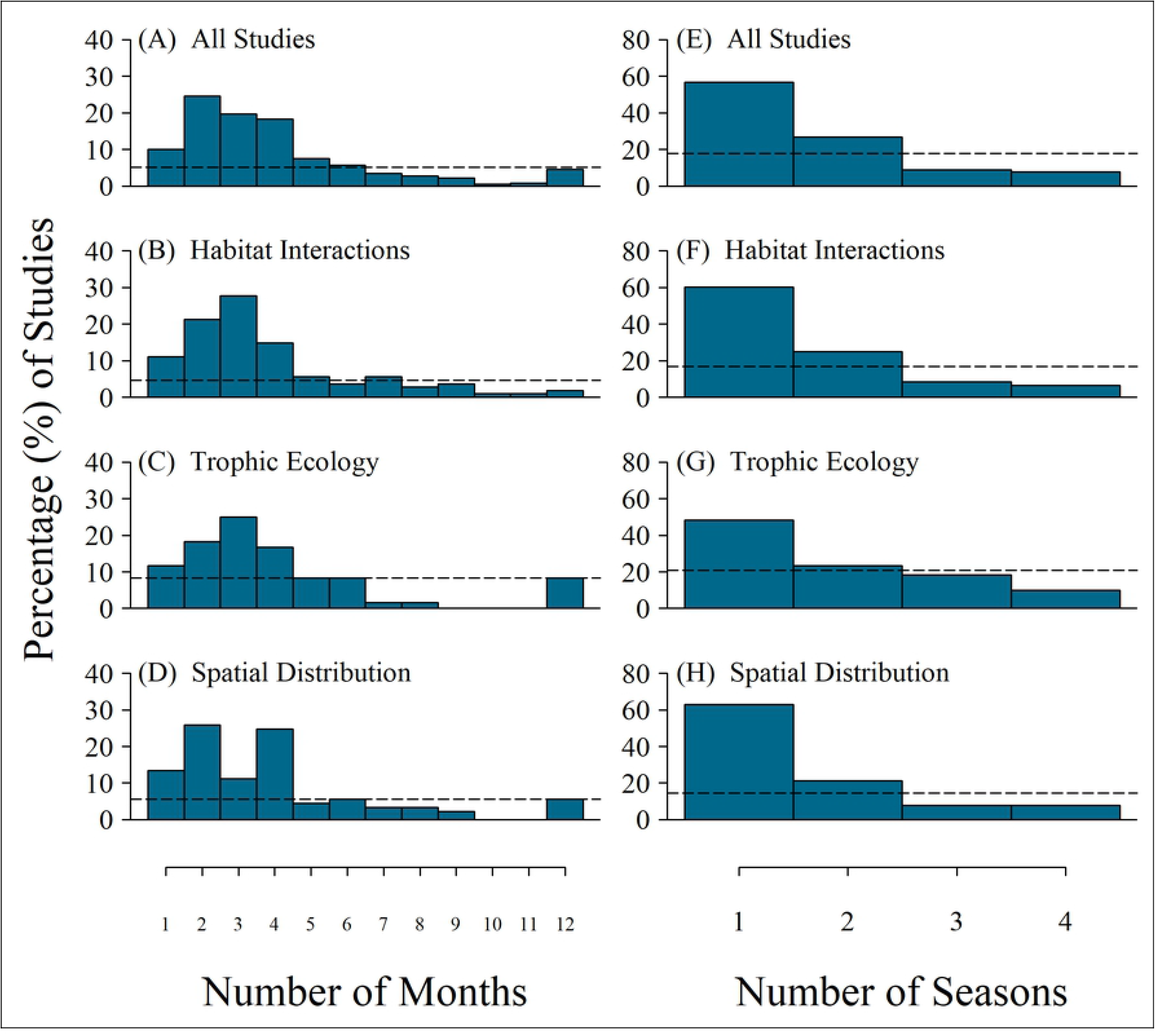
Temporal extent of juvenile salmon ecology studies. Left column: frequency of the number of months per calendar year (1-12) found in juvenile Pacific salmon and trout studies from 1988-2017 for (A) all studies (X2=670.07, p < 0.0001, n=371, median=5.1), (B) habitat studies (X2=173.55, p < 0.0001, n=108, median=4.6), (C) trophic ecology studies (X2=120.92, p < 0.0001, n=60, median=8.3), (D) spatial distribution studies (X2=173.01, p < 0.0001, n=89, median=5.1). Right column: frequency of the number of seasons per calendar year (1-4) found in juvenile Pacific salmon and trout studies from 1988-2017 for (E) all studies (X2=230.95, p < 0.0001, n=371, median=17.8), (F) habitat studies (X2=80.296, p < 0.0001, n=108, median=16.7), (G) trophic ecology studies (X2=19.6, p < 0.001, n=60, median=20.8), (H) spatial distribution studies (X2=72.573, p < 0.0001, n=89, median=14.6). The extent or duration was calculated by counting the total number of unique months (in a calendar year) that were included in each study and categorizing them by season as defined above. Data median is marked with a dashed horizontal line. Studies were only represented once at their greatest monthly extent and greatest seasonal extent.

### Seasonal temporal extent of studies

Across all ecological topics within juvenile *Oncorhynchus* spp. studies, we found that 48-63% of studies occurred during a single season while only 6-10% of studies encompassed field sampling from all four seasons (Fig 3). Only 43% of all studies collected data from multiple seasons and 73% of studies were shorter than 4 months. Again, there has been little change in the temporal extent of research efforts with the proportion of single-season studies remaining significantly higher than multi-season or year-round studies (Fig 4).

**Fig 4.**
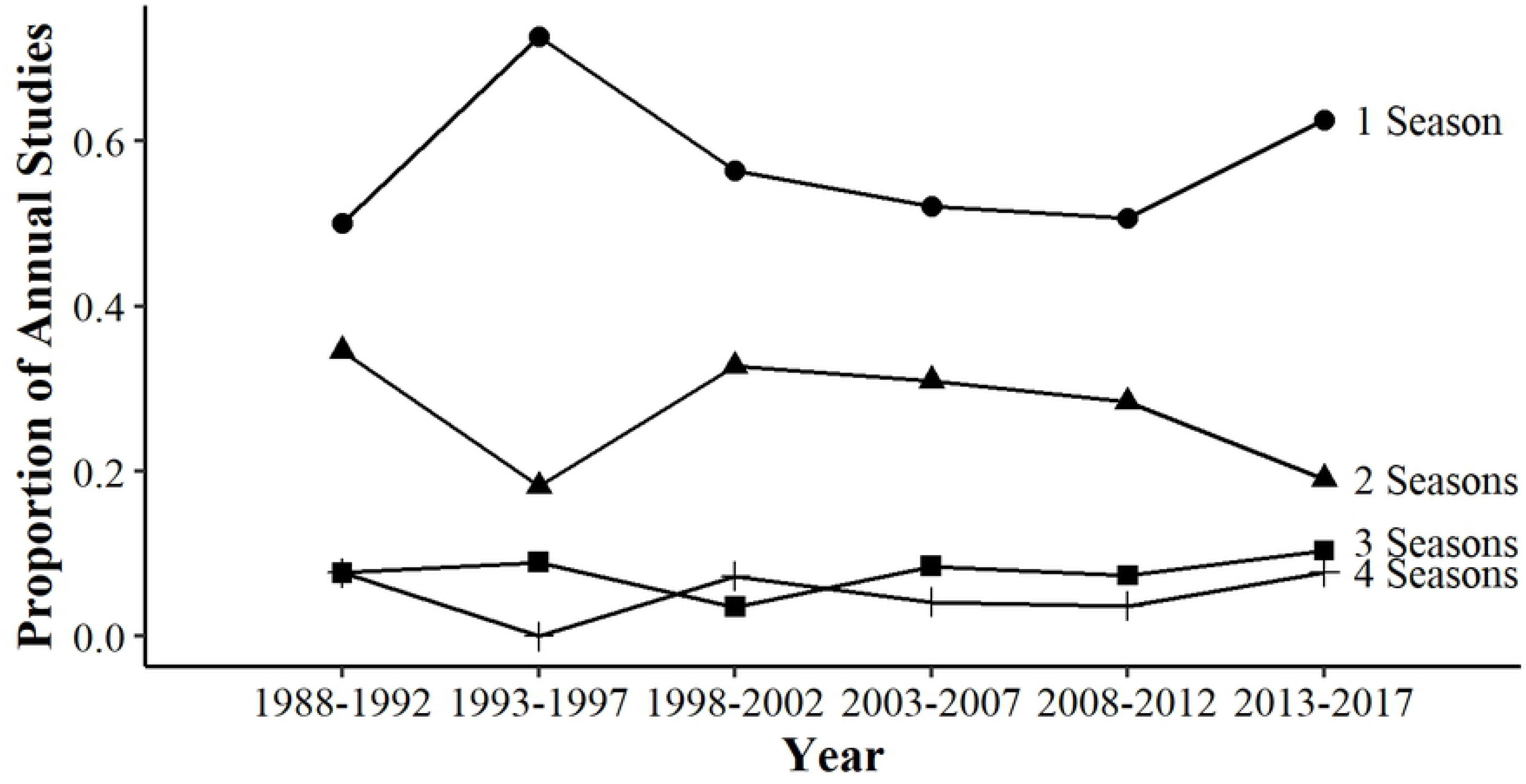
Seasonal study extent over time. Change in the proportional temporal extent (number of seasons included) of all studies published from 1988-2017 in 5-year increments.

### Riverscape studies

Analysis of riverscape studies across fish species revealed wider biases in temporal distribution at monthly and seasonal scales. The most frequently represented month was 8x more common than the least frequently represented month (Fig 5). January and February were the least represented months, while June, July, August, and September were most common. Summer encompassed 47% of all riverscape studies while only 11% of studies occurred during winter (Fig 5).

**Fig 5.**
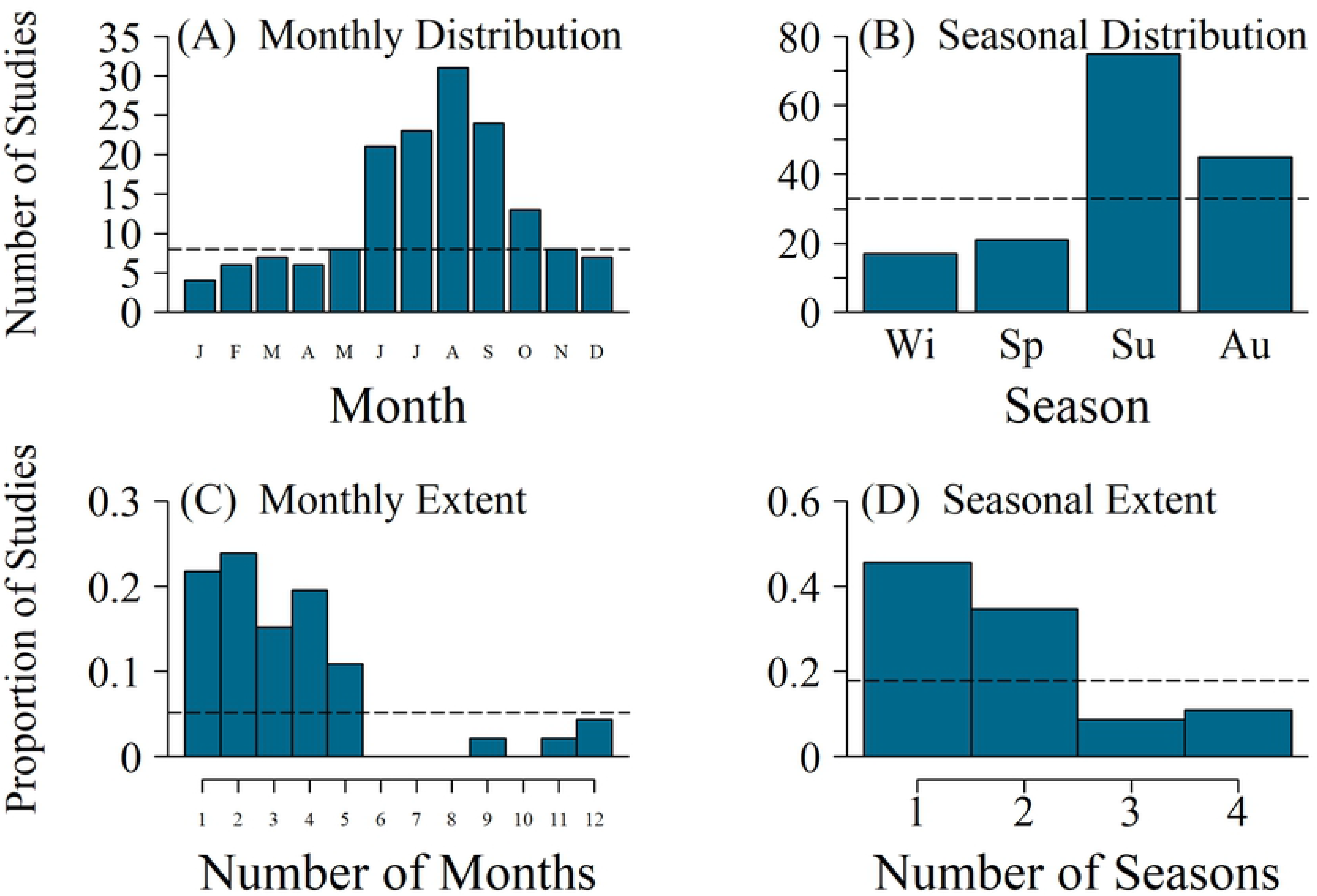
Distribution and extent of riverscape studies. (A) Monthly distribution (left to right: January to December) of sampling effort for spatially continuous “riverscape” studies involving all fish species from 1988-2017 (X2=69.089, p < 0.0001, n=158, median=8); (B) seasonal distribution of sampling effort for riverscape studies (X2=54.152, p < 0.0001, n=158, median=33); (C) frequency of the number of months per calendar year (1-12) found in riverscape studies (X2=97.038, p < 0.0001, n=46, median=3.3); (D) frequency of the number of seasons per calendar year (1-4) found in riverscape studies (X2=18.174, p < 0.001, n=46, median=22.83). The number of studies for each month or season was calculated using presence or absence of research during that time frame. Dashed horizontal lines are data median. Studies may occupy more than one month or season. Seasons were defined meteorologically, but as whole months. Summer is defined as the months June, July, and August; Autumn is defined as the months September, October, and November; Winter is defined as the months December, January, and February; Spring is defined as the months March, April, and May.

Monthly temporal extent was limited within riverscape studies as well. Spatially continuous studies were almost entirely conducted during a limited amount of time: 81% contained data from 4 months or less and only 4% of studies encompassed data from a full 12 months out of the year (Fig 5). Seasonal extent for riverscape studies was the one metric that was more representative than the ecological studies we examined: 46% of riverscape studies occurred during a single season, 35% occurred over two seasons, 9% occurred over three seasons, and 11% occurred during all four seasons (Fig 5).

## Discussion

In our review of 371 ecological juvenile *Oncorhynchus* spp. studies and 46 riverscape studies from the last 30 years, we observed strong biases in seasonal timing (distribution) and temporal extent. Within research topics where seasonality is particularly relevant, we observed the same general pattern of temporal bias; the period of summer was overrepresented in the study of fish-habitat interactions, trophic ecology, and spatial distribution. Below we discuss these temporal patterns of data collection and consider their potential causes and consequences.

### Bias in temporal distribution of studies

The most conspicuous pattern in the data was the lack of research during winter. For example, the month of December had less than one-quarter as many studies as that of June. Winter studies represented only 10-15% of total ecological research and 11% of riverscape studies. Winter may be tempting to overlook because it is generally a period of low biological activity in freshwater ecosystems. Winter is typically the coldest time of year, limiting the scope for growth and activity in aquatic poikilotherms. Further, winter is the darkest time of year, limiting primary productivity [19] and the foraging opportunity for visual predators [15]. Indeed, many stream-dwelling fishes tend to allocate energy to fat stores in anticipation of winter [20], suggesting it is generally a period of negative energy balance. However, this does not mean that understanding winter ecology is not critical. If fish rely on summer and fall fat stores to survive winter, then any food intake during winter helps to minimize the need to deplete those stores. Identifying winter foraging opportunities, trophic pathways, and habitat use could provide insights into how fish survive during this time of year [21]. For example, recent research exploring how environmental conditions influence fish interactions and movement has identified habitat not utilized outside of the winter months [22]. In many systems, winter survival is hypothesized to be a limiting factor to freshwater population productivity [23] and reducing winter mortality is often an objective of largescale restoration efforts [24]. Though juvenile salmonids may be less active in winter and not achieve substantial growth in many cases, there is evidence that winter fish growth may exceed growth observed during other seasons for some fish [25]. Understanding winter habitat use and foraging ecology could help improve our ability to increase overwinter survival.

The lack of winter research contrasted with the overabundance of summer studies. While emphasis on summer has benefits, such as an improved understanding of salmonid ecology during periods of climate stress, relying on summer-biased data could pose problems for conservation and management by violating assumptions of models. For example, species distribution models (SDM) are increasingly used in climate change adaptation and rely on the assumptions that a species occurs in all suitable habitats and that a species only occupies a portion of that suitable habitat due to constraining factors such as competition or predation [26]. Developing such models from temporally biased data would be valid only if the focal species were sedentary and their habitat use did not vary over time. However, it’s rarely possible to confirm that a species meets these criteria without having temporally representative data (i.e., you can’t dismiss the possibility of winter habitat shifts without data on winter habitat use). Using data from a limited period of time can cause SDMs to erroneously dismiss critically important habitat. For example, one study demonstrated that SDMs based on seasonally biased data failed to identify the habitats needed to support both hibernation and reproduction in bats [27]. Defining climate refugia for fish based on summer-biased data [28] could similarly leave out critical overwinter habitats if fish exhibit seasonal movements and require multiple habitat types to complete the annual cycle. While summer heat stress may be the most vivid threat of a warming world, climate change may also make winter more challenging by increasing maximum flows [29] or reducing ice cover [18]. The lack of winter studies in our analysis, and the emphasis on summer in both empirical studies and climate models [28], suggests that winter may be a blind spot for climate change adaptation work on Pacific salmon.

Our current classification system for longitudinal fish zonation is largely based on summer sampling [4]. While recent decades have seen an emphasis on more spatially representative fish sampling [30] and a movement towards multiscale analysis of spatial distributions [31], this work tends to not be temporally representative. For example, spatially continuous “riverscape” sampling has been transformative for our understanding of salmonid spatial distributions [6], yet our results confirm that virtually all of this work is conducted during summer or early autumn [32,33]. While longitudinal patterning is inherently relevant to lotic ecosystems (because they are linear networks), fish may also exhibit pronounced spatial patterning in lateral, and vertical dimensions [34]. In temperate regions of the Pacific salmon range, floodplains may only be connected and wetted during winter, so summer-biased sampling may hinder our ability to understand the significance of off-channel habitat use. Where summer and fall are the wet seasons (e.g., much of coastal Alaska), use of off-channel habitats may vary seasonally and require temporally extensive sampling to understand key dynamics. For example, the spatial patterning of juvenile coho salmon on a stream floodplain shifted over time, tracking shifts in water temperature [35] caused by fluctuating water levels. Use of temporary aquatic habitats by fish may be disproportionately important when they are available at the right place and time; however, research is lacking to capture this ephemeral aspect of fish ecology [36].

The distribution of juvenile salmonids among channel-unit scale habitat types [37] may also vary among months and seasons. For example, one study found that juvenile coho primarily occupied backwater pools in spring, main-channel pools in summer, and alcoves and beaver ponds in winter [38]. Distribution of juvenile salmonids in sub-habitats (e.g. riffles, pools, backchannel ponds) can also impact fish growth and fitness through energetic costs and benefits [39]. While fine-detail studies of fish distribution help identify quality salmonid habitat, our analysis demonstrates that this data implicitly favors summer habitat and devalues winter habitat.

### Bias in temporal extent of studies

While a bias against winter studies is seen in temporal distribution, a bias against full annual studies is seen in temporal extent. Ecological *Oncorhynchus* spp. studies examining all four meteorological seasons represented only 6-10% of total research. Research is heavily skewed toward shorter, single season studies: 73% of all studies capturing 4 months or less of data and 57% of studies focused on a single season in isolation. Within riverscape studies, 81% of research occurred during 4 or fewer calendar months. These patterns are similar to patterns observed in the phenology of mammal, bird, reptile and amphibian research [17]. While there is increasing recognition of the value of long-term study [40], this usually means having multiple years or decades of data collection. Our review shows that there is also a lack of temporal extent in terms of the annual cycle. Lacking extent at this timescale leads to two issues. First, we are likely to temporally extrapolate and draw conclusions based on a subset of the year (as discussed above) and second, we will often lack the ability to identify interactions between different time periods, or carry-over effects [17].

Carry-over effects from one life stage or season can have significant impacts on fitness and survival of individuals and populations in subsequent seasons or life stages [41]. As climate change and increasing water demands make summer more stressful for salmon in regions such as the western United States, there is a strong need to understand how conditions during spring and fall mediate the effects of summer stress on freshwater rearing capacity. The ability of fish to survive negative energy balance during harsh summer conditions should depend on their ability to store energy in spring and rebuild energy stores in fall. For example, over-winter survival of juvenile salmon is often positively associated with larger body size at the onset of autumn [42]. There is evidence that ephemeral food subsidy pulses, such as salmon eggs during the adult spawning season, can positively influence juvenile salmon growth rate and energy density as long as 6 months after this ephemeral resource pulse has disappeared [43]. Whether juvenile salmonids grow large enough to consume eggs depends on their emergence timing and early growth opportunities [44]. Thus, small increases in the growth of fry during spring may determine whether marine subsidies benefit parr during fall, influencing overwinter survival and the size of smolts the following spring, which relates to subsequent marine survival [45].

Sampling during multiple seasons is more likely to capture any carry-over effects that span pre-pulse, pulse, and post-pulse. Food availability, along with temperature, strongly affect fish growth rates with extreme variation in growth between seasons [25,46]. Quantifying fish growth and food resources at multiple points in time are essential to avoid bias in assumptions and to identify ephemeral trophic pathways that could be disproportionately important during that season or in subsequent seasons. Additionally, consequences of increased stress during one season can be observed in subsequent seasons through differences in fish growth, behavior, and survival [47,48]. Compensatory responses, such as growth rate and survival after a period of starvation, may also not be fully realized for many months [49,50]. The lack of full annual cycle research on Pacific salmon has likely hindered our ability to recognize inter-seasonal carry-over effects and compensatory responses, which may become increasingly important in the future.

A core concept in landscape ecology is that of habitat complementation and different patches of space functioning at different times (e.g., different life stages or seasons) [51,14]. The use of habitat by juvenile salmonids shifts 1) seasonally as river conditions such as temperature gradually change [38] 2) momentarily as a balance of energetic costs and benefits [52], 3) ontogenetically as resource needs change [2] and 4) ephemerally, such as during discrete events like floods or drought [14]. Without full annual studies, the effects of these stressors on fish (e.g. energetic costs, food availability, competition, predation) are poorly understood. Habitat restoration may be more successful if information is available to allow for targeting of the limiting life stage or limiting habitat in salmonid productivity [53]. Identification of these productivity limitations is hindered by two kinds of error: an assumption of limitation and an assumption of importance. First, the assumption that winter is limiting to juvenile salmonid survival is problematic because without more winter studies we cannot validate this assumption or understand the mechanisms behind winter mortality or winter vulnerability. Second, if we assume that summer is more important because significant growth occurs in the summer months, we assume that summer sampling can characterize spatial distribution and habitat use. This is problematic because it hinders the ability to identify limitations to juvenile salmonid survival outside of spring through fall. It is well-established that the challenges faced by stream-dwelling fishes in winter are vastly different [54]. To best protect the habitat supporting juvenile salmon and trout, more effort is needed to understand the importance of winter ecology.

### Considerations

The seasonal bias of research could potentially be a product of two human limitations: environmental challenges and allocation of scarce resources. First, the summer months generally present the least challenging environmental conditions for human access to salmon-bearing habitat, particularly in the Pacific Northwest where a significant amount of fish research takes place: low stream flow, warm temperatures, and minimal precipitation. Sampling fish in the winter months can be particularly challenging, as snow, ice, and high flow events limit safe access for researchers and lead to fish exhibiting behaviors that make them difficult to capture (e.g. winter concealment, nocturnality). Second, academic calendars create a seasonal bias towards summer field work by their very structure, allowing time for field work while classes are on break during summer. Field projects outside of academia also often follow a summer-intensive field season program due to the availability of field technicians who are often college students. Institutional hiring policies can further exaggerate these patterns. For example, at our institution students cannot work > 20 hours per week during non-summer months, and it costs ~30% more to hire seasonal assistants that are not students (due to the need for a temporary hiring agency). This makes non-summer field work considerably more expensive. Thus, a combination of environmental challenges, logistical hurdles, and institutional culture make field work more likely to happen in summer.

## Conclusion

In recent decades, stream ecology has strongly emphasized the need for more spatially comprehensive sampling of fish [6]; however, the focus on space has often come at the cost of time. Mapping the entire riverscape can reveal rich, multiscale patterns, but efforts typically fail to reveal how these patterns shift over time. Fish may not occupy every meter of space available to them, but they do live in every second of time. Furthermore, phenomena such as floodplain dynamics [1], seasonal movement [55], portfolio effects [56], resource waves [57], and thermoregulation [58] are driven by the interaction between spatial and temporal variation. We hope that our review encourages researchers to allocate more of their effort to understudied portions of the year, which likely hold valuable insights for conservation.

## Supporting information

**S1 Fig. PRISMA 2009 flow diagram for** *Oncorhynchus* **studies.**

(DOCX)

**S2 Fig. PRISMA 2009 flow diagram for riverscape studies.**

(DOCX)

**S3 Fig. PRISMA 2009 checklist.**

(DOCX)

**S1 Table. Articles included in***Oncorhynchus* **systematic review.**

(CSV)

**S2 Table. Articles included in riverscape systematic review.**

(CSV)

